# Increased Ovarian and Colorectal Cancer Cell Sensitivity to Platinum Drugs and innovative TS Inhibitors by Electroporation

**DOI:** 10.64898/2026.05.21.726835

**Authors:** Gaetano Marverti, Alice Belardo, Giada Mercanile, Daniele Aiello, Alberto Venturelli, Maria Paola Costi, Domenico D’Arca

**Affiliations:** Department of Biomedical, Metabolic and Neural Sciences, University of Modena and Reggio Emilia, 41125 Modena, Italy; Department of Life Sciences, University of Modena and Reggio Emilia, 41125 Modena, Italy

**Keywords:** Electrochemotherapy, Electroporation, Thymidylate Synthase, Cisplatin, Carboplatin, Oxaliplatin, Dimer disrupters

## Abstract

Ovarian and colorectal cancers have the highest incidence and mortality in the world, after breast cancer. Despite the initial response to Pt-drugs or 5-fluorouracil (5-FU), many cancer cells develop resistance to these drugs. For this reason, new therapeutic strategies represent an important medical need, in particular for drugs that are being studied in combination with methods that promote their entry into the cell. Among these strategies, electrochemotherapy (ECT), the combination of drugs with electroporation (EP), a physical method that uses high-frequency electrical pulses to create pores into which chemotherapy drugs can permeate, is gaining interest. In this study, we have evaluated the effect of ECT on the growth of both ovarian (A2780 and A2780/CP) and colorectal (HCT116) cancer cell lines using platinum derivatives (Cisplatin, Carboplatin and Oxaliplatin), as DNA alkylating agents, and human thymidylate synthase (hTS) inhibitors, both traditional (5FU) and novel TS destabilizers (compounds E3 and E7). To this aim, synergism quotient-like analysis to determine whether electroporation gives an advantage in terms of cytotoxicity was applied to the relative IC20 and IC50 concentrations of each drug. Results showed that two of the three Pt-drugs have greater efficacy when combined with EP. 5-FU and the new TS inhibitors E3 and E7 also take advantage of ECT because EP increases drug uptake into the cell, even in resistant cells. In conclusion, ECT appears to be a viable strategy to obviate the problem of resistance in ovarian and colorectal cancers, to deliver compounds inside cells overcoming uptake limits, especially for the low lipophilic compounds whose cytotoxic efficacy is hampered by the obstacle of biological membranes.

## Introduction

For Ovarian cancer (OC) is a complex, heterogeneous malignancy that remains a leading cause of mortality among gynaecological neoplasms. The therapeutic cornerstone involves cytoreductive surgery to minimize anatomical disease burden, followed by systemic chemotherapy. Primary treatment typically consists of platinum-based compounds specifically cisplatin or carboplatin combined with taxanes, such as paclitaxel (Tavares et al., 2024).

Cisplatin (cDDP) remains one of the most potent agents in the OC armamentarium. Its efficacy stems from its alkylating activity, which allows it to form DNA adducts that disrupt cellular replication.

While cDDP primarily enters both malignant and healthy cells via passive diffusion, its clinical utility is often limited by significant nephrotoxicity and ototoxicity. Consequently, carboplatin (CarboPt) was developed as a less toxic second-generation analogue; while it shares a similar mechanism of action, its improved safety profile permits higher dosing. In cases of recurrence, the therapeutic landscape expands to include agents such as topotecan, liposomal doxorubicin, gemcitabine, and trabectedin, frequently administered alongside targeted therapies like bevacizumab (anti-VEGF) and PARP inhibitors (Ferreira et al., 2016; Tavares et al., 2024). In ovarian cancer treatment, CarboPt is typically combined with paclitaxel; however, it demonstrates lower efficacy than cisplatin in treating testicular, bladder, and head and neck cancers.

Beyond passive diffusion, the intracellular accumulation of these platinum-based drugs is mediated by Copper Transporter 1 (CTR1), a transmembrane protein essential for copper homeostasis. The downregulation of CTR1 is a significant factor in reduced drug uptake and the subsequent development of platinum resistance. Conversely, the copper-extruding P-type ATPases, ATP7A and ATP7B, facilitate drug efflux: ATP7A sequesters cisplatin within intracellular vesicles, while ATP7B promotes its systemic release from the cell. In patients with ovarian or colorectal cancers, high expression levels of these transporters are clinical indicators of poor drug response and a worse overall prognosis (Gheorghe-Cetean et al., 2017; Dilruba and G. V. Kalayda, 2016).

Colorectal cancer (CRC) remains a global health challenge, ranking as the second leading cause of cancer-related incidence and mortality across both sexes. It accounts for approximately 10% of all annual cancer diagnoses and deaths worldwide. While surgical resection remains the gold standard for treatment, particularly in the early stages, more advanced or extensive disease requires a multimodal approach. In these cases, patients undergo systemic chemotherapy or radiotherapy in neoadjuvant (pre-operative) or adjuvant (post-operative) settings to improve resectability and mitigate the risk of recurrence.

The pharmacological arsenal for CRC primarily includes: Platinum derivatives, in particular Oxaliplatin (OxaPt); Pyrimidine analogues: 5-fluorouracil (5-FU) and its oral prodrug, Capecitabine; Topoisomerase I inhibitors: Irinotecan. Current first-line standards of care favour combination regimens over monotherapy due to their superior clinical efficacy. These standardized protocols include: FOLFOX (Folinic acid, 5-FU, and Oxaliplatin); FOLFIRI (Folinic acid, 5-FU and Irinotecan) and FOLFOXIRI, a triplet regimen combining Folinic acid, 5-FU, Oxaliplatin and Irinotecan (Dekker et al., 2019; Kumar et al., 2023).

OxaPt was developed to overcome resistance against cDDP and CarboPt since it forms a hydrophobic complex, bulky, hydrophobic DNA adducts that cause greater inhibition of DNA synthesis than its predecessors (Gheorghe-Cetean et al., 2017). The distinct lipophilicity of this third-generation compound enables it to utilize alternative transport pathways for cellular entry, specifically the organic cation transporters (OCT). The overexpression of OCT in CRC cells likely explains the unique efficacy of OxaPt in this malignancy when administered alongside 5-FU and folinic acid. Currently, OxaPt is the clinical standard for colorectal, gastric, and ovarian cancers, with ongoing trials investigating its utility in pancreatic, breast, and lung malignancies.

A primary pharmacological target in these tumors is the overexpression of thymidylate synthase (TS), a rate-limiting enzyme in the folate-dependent de novo pyrimidine synthesis pathway. TS catalyzes the reductive methylation of deoxyuridine monophosphate (dUMP) to deoxythymidine monophosphate (dTMP), utilizing 5,10-methylenetetrahydrofolate (mTHF) as a methyl donor.

Thymidylate synthase (TS) is a highly conserved homodimer, composed of 35 kDa subunits, that serves both catalytic and regulatory functions. Beyond its role in nucleotide synthesis, TS exerts negative feedback on its own expression by binding to its own mRNA, thereby inhibiting translational processing.

In the context of cancer, uncontrolled cellular proliferation demands an accelerated rate of nucleotide synthesis to sustain rapid DNA replication. Consequently, TS overexpression is frequently observed in malignancies such as ovarian and colorectal cancers. This upregulation is a primary driver of chemoresistance and positions TS as a critical target for therapeutic intervention.

5-FU is a pyrimidine antimetabolite that undergoes intracellular conversion into several active metabolites, most notably fluorodeoxyuridine monophosphate (FdUMP). The cytotoxic effect of 5-FU is achieved through two primary pathways, i. enzymatic Inhibition in which FdUMP acts as a fraudulent substrate, competing with natural dUMP for the nucleotide-binding site of TS. It forms a stable, covalent ternary complex with the enzyme and the cofactor mTHF. This locks the enzyme in an inactive state, depleting dTMP pools and causing a catastrophic imbalance in nucleotide levels that halts DNA synthesis. ii. Genotoxic incorporation in which a second metabolite, FdUTP, is erroneously incorporated into DNA strands. This misincorporation triggers lethal DNA damage and promotes apoptosis (Costi et al., 2005; Chu et al., 1991; Taddia et al., 2015).

For decades, 5-FU has been a cornerstone in the treatment of both early-stage and advanced colorectal cancer. To maximize its clinical efficacy, it is co-administered with Leucovorin (folinic acid). Leucovorin acts as a biochemical modulator by expanding the intracellular pool of methylenetetrahydrofolate (mTHF), which is essential for the formation and stabilization of the inactive FdUMP-TS-mTHF ternary complex. Without sufficient mTHF, 5-FU alone often fails to sustain enzyme inhibition, leading to the translational derepression of TS. This results in compensatory enzyme overexpression, a primary driver of 5-FU resistance.

A major challenge in contemporary oncology is developing novel strategies to circumvent this resistance. A promising therapeutic approach shifts focus from the active site to the enzyme’s quaternary structure. This strategy utilizes small-molecule TS inhibitors designed to bind at the dimer interface.

By disrupting the monomer-dimer equilibrium of this obligate homodimer, these molecules favor the formation of the inactive monomer. Unlike traditional inhibitors, these “dimer disrupters” (Ddis) offer a multi-modal effect such as enzymatic inhibition that prevents the assembly of the active catalytic site; protein degradation by destabilizing the homodimeric structure, they promote the intracellular degradation of the protein; reduced TS accumulation by preventing the surge in TS levels typically seen with active-site inhibitors, effectively neutralizing the mechanism of translational derepression (Costantino et al., 2022).

In this study, we evaluated two candidate Ddis compounds, LC1236 (**E3**) and LC1343 (**E7**), which exhibit distinct pharmacokinetic profiles. While E3 demonstrates superior inhibitory potency against the recombinant enzyme in vitro, its biological activity is hampered by poor transmembrane permeability. In contrast, **E7** possesses higher lipophilicity, allowing for efficient autonomous cell entry and robust intracellular activity. The mechanism of **E7** is particularly innovative; it accelerates the proteasomal degradation of TS, thereby decoupling enzymatic inhibition from the typical compensatory surge in protein expression that leads to resistance (Costantino et al., 2022).

Despite their potential, the clinical application of these compounds is often limited by suboptimal pharmacokinetics, characterized by low aqueous solubility and poor systemic absorption. Furthermore, cancer cells frequently employ multifactorial chemoresistance strategies, such as the downregulation of influx transporters and the upregulation of efflux pumps, which diminish the intracellular concentrations of 5-FU, platinum-based drugs, and novel inhibitors alike.

To circumvent these barriers, we have explored strategies to enhance drug internalization and maximize therapeutic efficacy. Recent findings demonstrate that the potency of both 5-FU and these innovative TS Ddis can be significantly enhanced by co-administration with specific inhibitors of ATP-binding cassette (ABC) transporter proteins to prevent drug extrusion; by utilizing an unconventional peptide-based delivery system, the SAINT-Protein transfection agent (Synvolux Products & Therapeutics™, Leiden, The Netherlands) to enhance drug influx. This delivery platform facilitates the intracellular transport of poorly permeable molecules, effectively enabling less lipophilic compounds like E3 to bypass the plasma membrane barrier (Marverti et al., 2024).

In the present study, we sought to bypass restricted cellular uptake and achieve therapeutic intracellular drug concentrations capable of neutralizing established resistance mechanisms. To this aim, we utilized electroporation (EP) to potentiate the activity of these compounds. This clinical integration of electric fields and cytotoxic agents is termed electrochemotherapy (ECT), a strategy aimed at optimizing treatment efficacy for ovarian and colorectal malignancies.

EP is a biophysical phenomenon triggered by the application of external electric pulses, which induce a transient increase in transmembrane potential. Once this potential exceeds a specific threshold, it leads to the stochastic formation of aqueous nanopores in the lipid bilayer. These pores act as conduits for water, ions, and large or charged molecules that are otherwise membrane-impermeant. When the electric field parameters are carefully calibrated, these pores close within seconds to minutes, a process known as reversible electroporation. This temporary increase in membrane permeability facilitates a massive influx of chemotherapeutic agents directly into the cytosol of tumor cells. By significantly enhancing intracellular accumulation, ECT increases cytotoxicity while allowing for a reduction in total systemic dosage, thereby mitigating treatment-associated side effect (Łapińska et al., 2022; Kumar et al., 2019). Among the first drugs tested in combination with EP, those that gave better results were bleomycin (BLM) and cisplatin (cDDP), whose toxicity increases on average of 1000- and 80-fold, respectively (H. Falk Hansen et al., 2020; Perrone et al 1993). 5FU and cDDP have been also tested with EP in ovarian cancer cells (Saczko et al., 2014).

The primary objective of this study was to determine whether EP confers a synergistic cytotoxicity advantage to a panel of compounds with diverse pharmacokinetic profiles. This panel included established clinical standards cDDP, CarboPt, OxaPt and 5-FU alongside the novel allosteric TS inhibitors, **E3** and **E7**.

To evaluate the potential of ECT to circumvent multidrug resistance (MDR), we utilized cell models representing two of the most lethal malignancies: ovarian and colorectal cancer. Specifically, the study employed A2780 (Ovarian Cancer), a sensitive parental line used as a baseline for comparison; A2780/CP (Ovarian Cancer), an acquired-resistance model, characterized by its resistance to cDDP and cross-resistance to 5-FU and HCT116 (Colorectal Cancer), a model chosen for its innate (intrinsic) low sensitivity to 5-FU. By comparing the efficacy of these agents with and without EP across these sensitive and resistant phenotypes, we aimed to establish a more effective therapeutic strategy for overcoming pharmacological barriers in refractory tumors.

## 2. Materials and Methods

### 2.1. Drugs and chemicals

The following anticancer drugs were used: cisplatin (Sigma), carboplatin (Sigma), oxaliplatin (Sigma), and 5-fluorouracil (5FU) (Sigma). Drugs that were received as solids were dissolved in sterile 0.9% bacteriostatic saline. Also, two investigational compounds were tested: LC1236 (**E3**) (2-Phenyl-2-(4-nitrophenylthio)-N-(4-carboxyphenyl) carboxylic acid) and LC1343 (**E7**) (phenyl-2-(4-nitrophenylthio)-N-(6-methanesulfonylbenzothiazol-2-yl)acetamide), synthesized in the Drug Discovery and Biotechnology Lab (P.I. Maria P. Costi, Unimore) following the methods previously reported. The compounds were used as racemic mixtures (SI). **E3** and **E7** were chosen because of their different pharmacokinetics and subsequent pharmacodynamics. **E3** shows limited uptake through the plasma membrane, while **E7** can be easily internalized.

### 2.2. Cell cultures

HCT116 (Colorectal Adenocarcinoma cell line) was cultured in Dulbecco Modified Eagle Medium (DMEM), supplemented with 10% heat-inactivated fetal bovine serum and 1% Pen/Strep. The HCT116 cell line was purchased from ATCC (American Type Culture Collection, Manassas, VA, USA). A2780 and A2780/CP, ovarian carcinoma cell lines, were cultured in RPMI 1640 medium containing 10% heat-inactivated fetal bovine serum, 1% Pen/Strep. The A2780/CP cells are about 10-fold resistant to cisplatin and derived from the parent A2780 cell line. The A2780 and A2780/CP cell lines are a generous gift of Prof. E. M. Berns, Department of Medical Oncology, Erasmus MC Cancer Institute, Rotterdam, The Netherlands, and were purchased from the European Collection of cell cultures, ECACC via Sigma (Beaufort et al., 2014).

Cells used for experimental purposes were detached using trypsin-EDTA (0.05% trypsin-0.53 mM EDTA; Mediatech), collected by centrifugation at 900 rpm for 5 min. Cells were then resuspended in the electroporation buffer (RPMI 1640, 10× diluted 1:10 without glutamine and serum) and enumerated using a hemacytometer (Fischer Scientific, Pittsburgh, PA). Cells with a viability greater than 95%, as determined by the Trypan blue dye method (0.4% Trypan blue; Sigma), were used for all experimental purposes.

### 2.3. Crystal violet assay for cell survival

After electroporation alone or electroporation in the presence of a drug, aliquots containing 0.4×10^6^ cells (in 80 μL) were seeded into 6-well tissue culture plates (Falcon, Lincoln Park, NJ). Wells were then filled with 3 ml of the appropriate media. Plates were incubated at 37°C for 24 h for the case of electroporation alone and for 72 h when cells were porated in the presence of anticancer drugs. From the plate, the culture medium was removed, and the monolayer of cells was fixed with methanol and then stained with a solution of 0.2% violet crystal in 20% methanol for at least 60 minutes (Feoktistova et al., 2016; Marverti et al., 2019)

At the end of the treatment, cellular staining is obtained as crystal violet can bind to DNA and proteins. The excess dye was removed, which was not incorporated, and the cells were left to dry overnight. The built-in dye was solubilized with a solution of acid isopropanol (1 N HCl: 2-propanol, 1:10), adding 2 mL to each well. The dye well concentration is proportional to the number of cells. The absorbance was determined by spectrophotometric analysis at a wavelength of 540 nm using a 24-well plate reader (Tecan GeniusPro). Finally, the percentage of cytotoxicity was calculated by comparing the absorbance of cells exposed to compounds and compounds + EP with the absorbance of cells not exposed, i.e., negative control.

### 2.4. Cytotoxicity of drugs without electroporation

On the first day, 5x10^4^ cells were seeded per well on a 24-well plate and left for one day in an incubator at 37°C and humidified 5% CO2. On the second day (24 h after cell seeding), drugs were mixed with cells at different concentrations (from 5 to 100 μM) and were incubated for 48 h or 72 h. After the indicated time, the Crystal Violet dye assay was performed (Feoktistova et al., 2016; Marverti et al., 2019). The same procedure was performed for all tested compounds. From the dose-response curves obtained, the IC20 and IC50 of the tested compounds were extrapolated. The IC50 and IC20 are defined as the concentration causing 50% and 20%, respectively, growth inhibition in treated cells when compared to control cells after the indicated drug exposure times.

### 2.5. Electroporation apparatus

The Genedrive electroporator (IGEA, Carpi, MO, Italy) was used to generate rectangular direct current pulses (Perrone et al., 1993). Electroporation-treated samples received eight 100 μs pulses at 5 KHz. Electric field strengths were varied by adjusting the output voltage of the generator. Cell suspensions were placed into standard 2 mm gap electroporation cuvettes (Bio-Rad Laboratories S.r.l.). The cuvettes were constructed with parallel aluminium plates which served as electrodes. Aliquots containing 0,4×10^6^ cells (in 80 μl) were placed into the cuvettes and electric pulses of a specific electric field strength were applied using the generator. The Genedrive electroporator was used to monitor duration, voltage and current associated with each delivered pulse.

### 2.6. Cell survival after electrical treatment of EP

Aliquots of 0.4×106 cells (in 80 μl) were placed into separate electroporation cuvettes. Cells in each cuvette were electroporated at a specific electric field strength. This sample procedure was conducted for electric field strengths of 500, 750, 900, 1100, and 1250 V/cm. Ten minutes after electric field exposure, aliquots of 0.4×106cells were plated from each cuvette into a 6-well tissue culture plate. The plate was incubated for 24 h at 37 °C, then a crystal violet dye assay was performed. Experiments were run in triplicate, and the electric field strength which maintained about 75% survival was determined by examining mean plots of survival versus electric field strength.

### 2.7. Synergy analysis of drug combination by synergism quotient

The effects of drug combination were quantified by the synergism quotient (SQ) (Y. S. Cho and Y. S. Cho-Chung, 2003; G. Marverti et al., 2020). SQ was defined as the net growth inhibitory effect of the drug combination divided by the sum of the net individual drug effects on growth inhibition. A quotient of > 1 indicates a synergistic effect, between 0.9 and 1.1 indicates an additive effect, while a quotient of < 1 indicates an antagonistic effect.

### 2.8. Statistical analyses

All values are reported as the mean ± standard deviation (SD), unless otherwise indicated. Statistical significance was determined using a two-tailed Student’s *t*-test performed in Microsoft Excel. Differences were considered statistically significant at *P <0.05 or **P <0.01; *** p < 0.001.

## 3. Results

### 3.1. Identification of electrical parameters to obtain reversible electroporation by propidium iodide

First, we performed experiments to determine the optimal electroporation (EP) parameters to be used in subsequent experiments with chemotherapeutic drugs. Optimal electroporation parameters were defined as those that achieved the highest cell membrane permeability and the highest cell survival. We started by evaluating the increase in permeability caused by different voltages. For this purpose, Propidium Iodide (PI), a fluorescent dye which cannot permeate the cell membrane, was used in combination with electroporation to monitor the enhancement of permeability. As a measure of permeability, we used the fluorescence signal emitted upon the binding of PI to cellular DNA.

In **Figure 1**, the percentage of PI which enters the cells is depicted. The permeability of the membrane increases with the strength of the applied electric field. The highest uptake was achieved at 1100 V/cm and at 1250 V/cm (**Figure 1, panels C and D**). Then, cell survival was measured after electroporation alone, using the same voltages as in the previous experiment. The electric field strength which maintained about 75% survival was determined by examining mean plots of survival in different cell lines versus electric field strength. The electric pulses at 900 V/cm lead to about 75% of survival, 1100 V/cm induces about 72% of survival, and 1250 V/cm gives about 55% of survival. 900 V/cm and 1100 V/cm intensities represents the electric field strengths that maintained ∼75% of survival. For the following ECT experiments, 1100 V/cm was chosen because it achieved higher uptake than 900 V/cm, as shown in **Figure 1**, but a comparable effect on cell survival.

**Figure 1.**
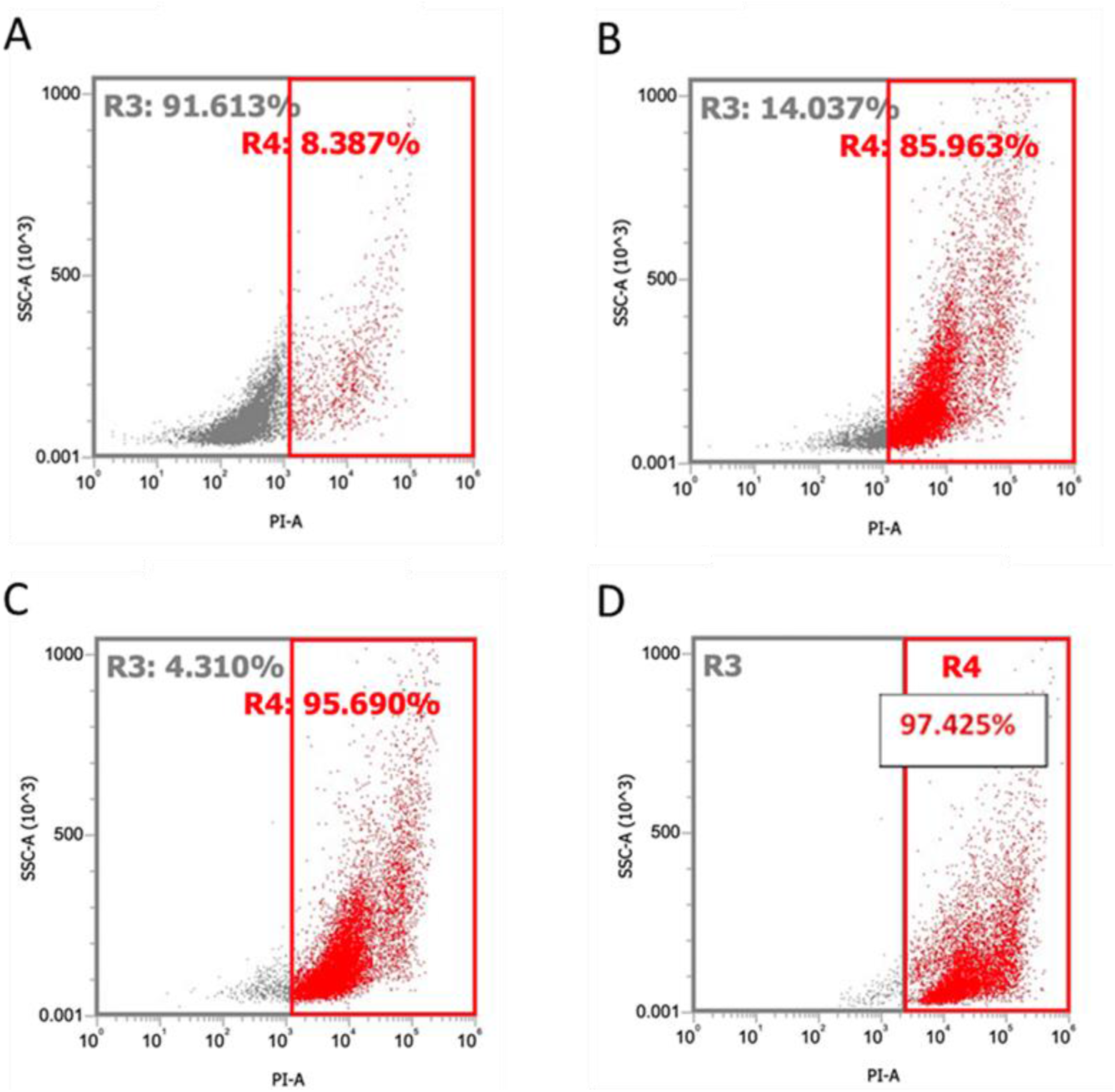
Electroporation with Propidium Iodide: (A) control (no electroporation); (B) 900 V/cm; (C) 1100 V/cm; (D) 1250 V/cm.

### 3.2. Cytotoxicity of drugs without electroporation for IC20 and IC50 determination

Before starting the EP experiments, a critical step was the exact determination of the inhibitory concentrations, specifically the IC20 and IC50 values. These metrics serve as vital benchmarks, defining the drug concentrations at which 20% and 50% of cell growth or viability is inhibited, respectively.

The focus of this initial investigation centred on understanding the sensitivity of several key cancer cell lines. Two OC cell lines were chosen for their distinct characteristics: A2780, representing a drug-sensitive parental line, and A2780/CP, its acquired cisplatin-resistant counterpart. This pairing allowed for a comparative analysis of drug efficacy against both sensitive and acquired resistance phenotypes, providing insights into potential therapeutic strategies. Complementing these, the CRC cell line HCT116 was also included, broadening the scope of the study and allowing for the assessment of drug activity across different cancer types.

These selected cell lines were then subjected to a panel of therapeutic agents. The established platinum-based compounds, were tested to provide a reference point for drug activity. Alongside these conventional agents, two novel and promising compounds, **E3** and **E7**, were introduced. These compounds represent potential new avenues in cancer therapy, and their initial characterization in these cell lines was crucial. Both compounds were tested in previous experiments. Compound **E7** was the most active one and showed the capacity to be internalized. Instead, compound **E3** was not able to cross the cell membrane (Marverti et al., 2024). To obtain the precise IC20 and IC50 values, a series of dose-response curves was obtained for each drug-cell line pair. This involved exposing the cells to a range of drug concentrations, from very low, sub-inhibitory doses to concentrations expected to induce significant cell death. The cellular response was then quantified after two distinct incubation periods: 48 hours and 72 hours. This dual time-point analysis was critical for understanding not only the immediate effects of the drugs but also their sustained impact over a slightly longer duration, offering a more comprehensive picture of their cytotoxic profiles. These drug concentrations would then serve as the starting points for the subsequent EP experiments, ensuring that the EP parameters could be optimized in conjunction with biologically relevant drug doses.

**Figure 2** shows that OxaPt was the most active Pt-drug against all cell lines. Among the TS inhibitors, **E3** was the least effective drug without reaching an IC50 value within 100 µM. On the contrary, both 5-FU and **E7** were more active in all three cell lines, showing IC50 values below 25 µM (**Figure 3**). From the dose-response curves are inferred the IC20 and IC50 values of all drugs tested in the three cell lines and reported in **Figure 3**.

In greater detail, both ovarian cancer cell lines are much more sensitive to OxaPt and cDDP and much less, as expected, to CarboPt. In fact, the 72-hour IC50 values of the first two drugs are less than 15 μM in sensitive cells and 40 μM in resistant cell lines. In contrast, for CarboPt, values of 80 μM and above were found (**Figs. 2, 3**).

**Figure 2.**
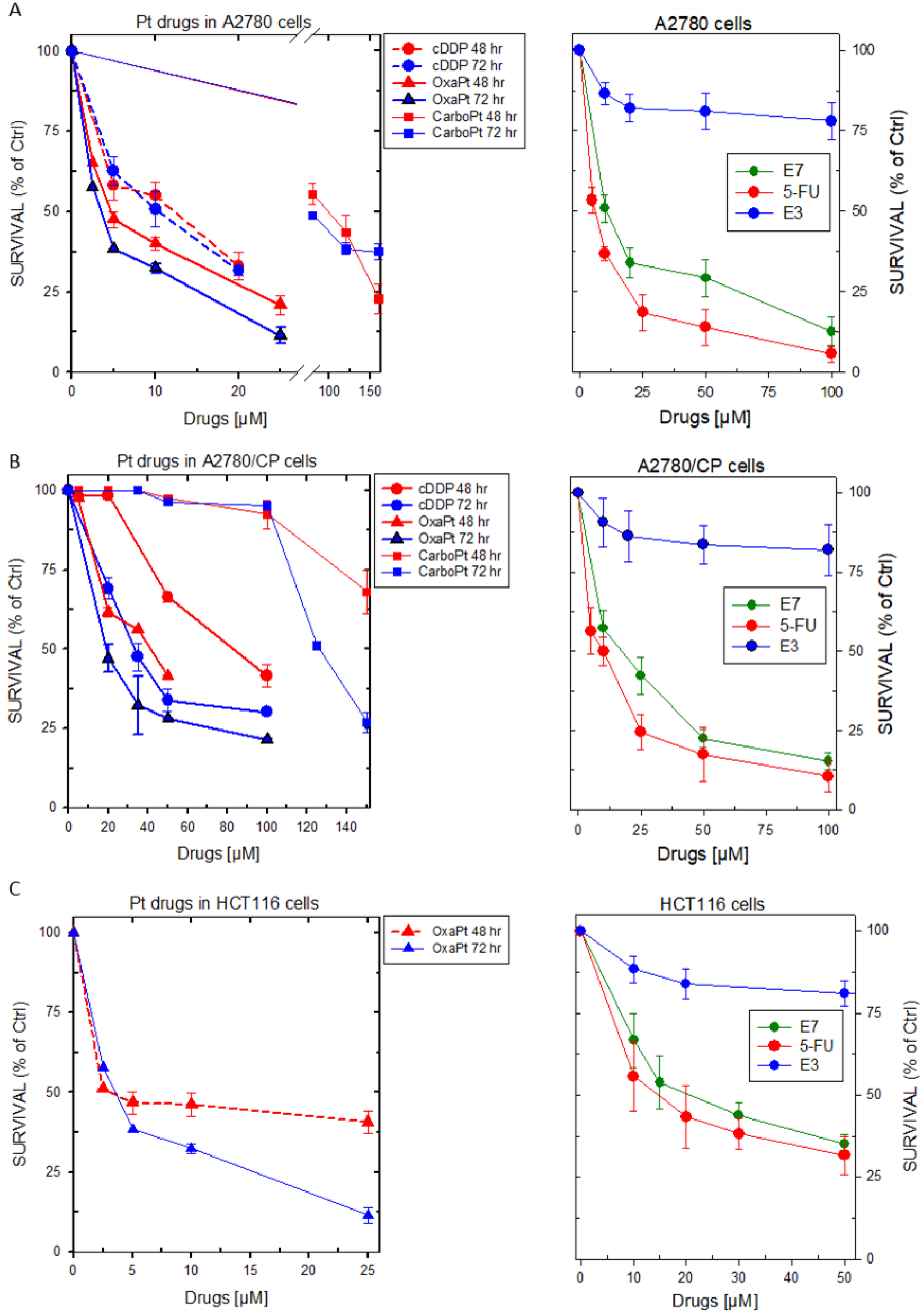
Dose-response curves for tested Pt-drugs (left panels), 5-FU, **E7** and **E3** compounds (right panels) in A2780 (A), A2780/CP (B) and HCT116 (C) cell lines. Cell survival rates were obtained after exposure to the indicated drug concentrations for 48 hr and 72 hr. Symbols are the means of three separate experiments performed in duplicate. Error bars, S.D.

**Figure 3.**
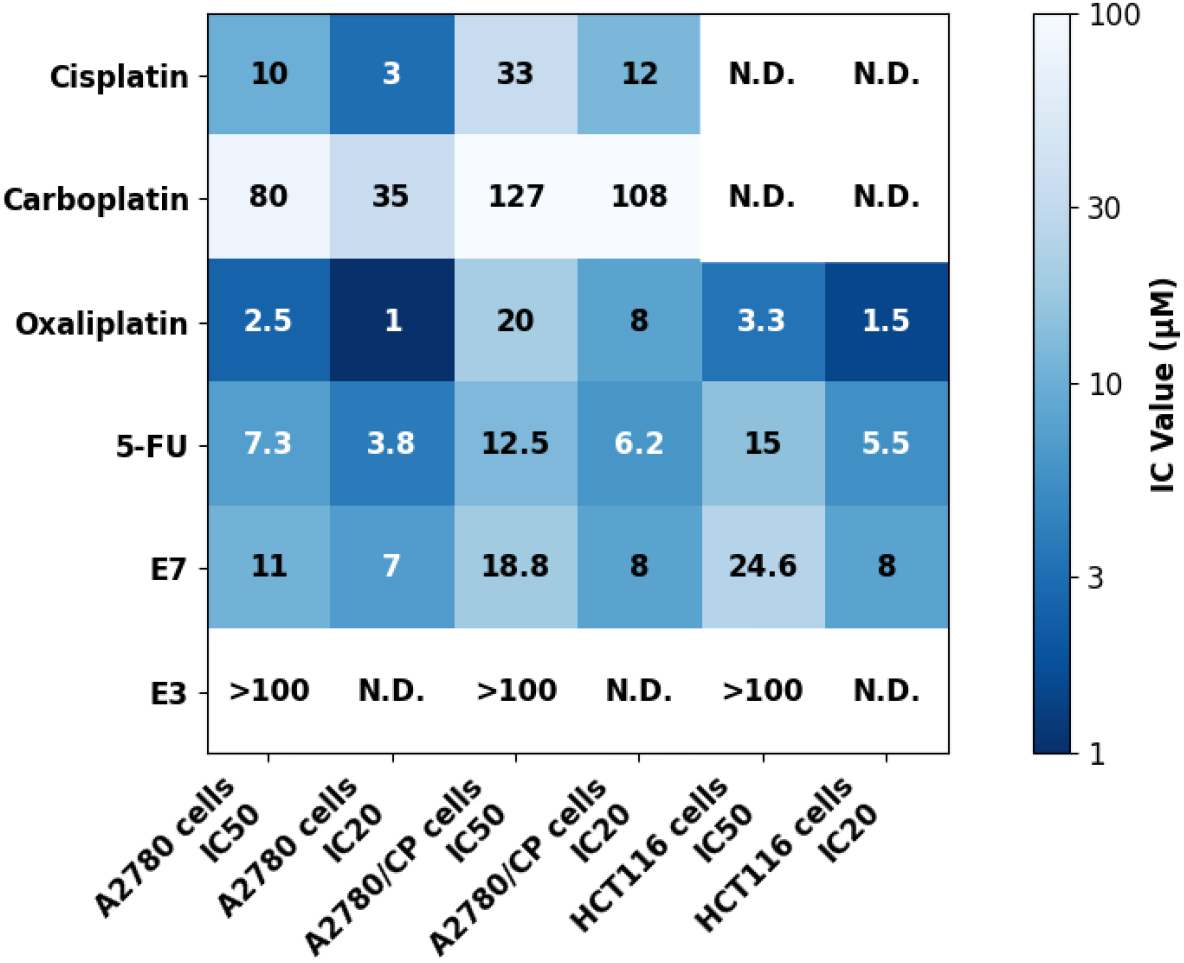
Heat-map representation of IC20 (μM) and IC50 (μM) values (20% and 50% cell growth inhibitory concentration) for the indicated compounds after 72 hr treatment of A2780 and A2780/CP OC cell lines and HCT116 CRC cell line. The values are the mean of at least three experiments. N.D., not determined. In the colour scale, darker colours mean lower IC50 values and greater drug efficacy.

Due to its lower lipophilicity, **E3** did not reach the IC50 values up to 100 μM, while **E7**, more lipophilic and with a higher logP (Costantino et al., 2022; Marverti et al., 2024), showed IC50 values lower than 20 μM and therefore an efficacy comparable to the traditional drug 5FU (**Figure 2**, right panels). The resistance phenotype of A2780/CP cells affected the effectiveness of most drugs, increasing the IC50 values in these cells in comparison to the sensitive A2780 cells.

Only the Pt-drug of choice for CRC, OxaPt, was tested in the CRC HCT116 cells, and in fact, it showed IC50 values lower than 5 μM. Even in this cell line, **E7** was almost as active as 5FU, while **E3** confirmed its poor cytotoxicity (**Figure 2C**).

### 3.3. Increased OC and CRC cell line sensitivity to Pt-drugs and to TS inhibitors by electroporation

For the ECT experiments, all drugs were used at the previously calculated IC20 and IC50 concentrations on the three cell lines and at the best EP conditions of higher uptake and the lowest cytotoxicity, that is 1100 V/cm. We started to evaluate ECT effects on the cisplatin-sensitive OC A2780 cells. As shown in **Figure 4A**, EP significantly increases the cytotoxicity of cDDP at both concentrations, since both IC20 and IC50 cDDP efficacy increased by about 45% and 30%, respectively, in comparison to cisplatin alone. The IC20 value of CarboPt alone caused approximately 20% cell growth inhibition, while the same concentration in the presence of EP caused 55% cell growth inhibition. A comparable and significant enhancement of cell killing was also observed for the IC50 concentration. Regarding OxaPt, the cell growth inhibition increased with ECT, but it was not sufficiently significant versus drug alone treatment.

**Figure 4.**
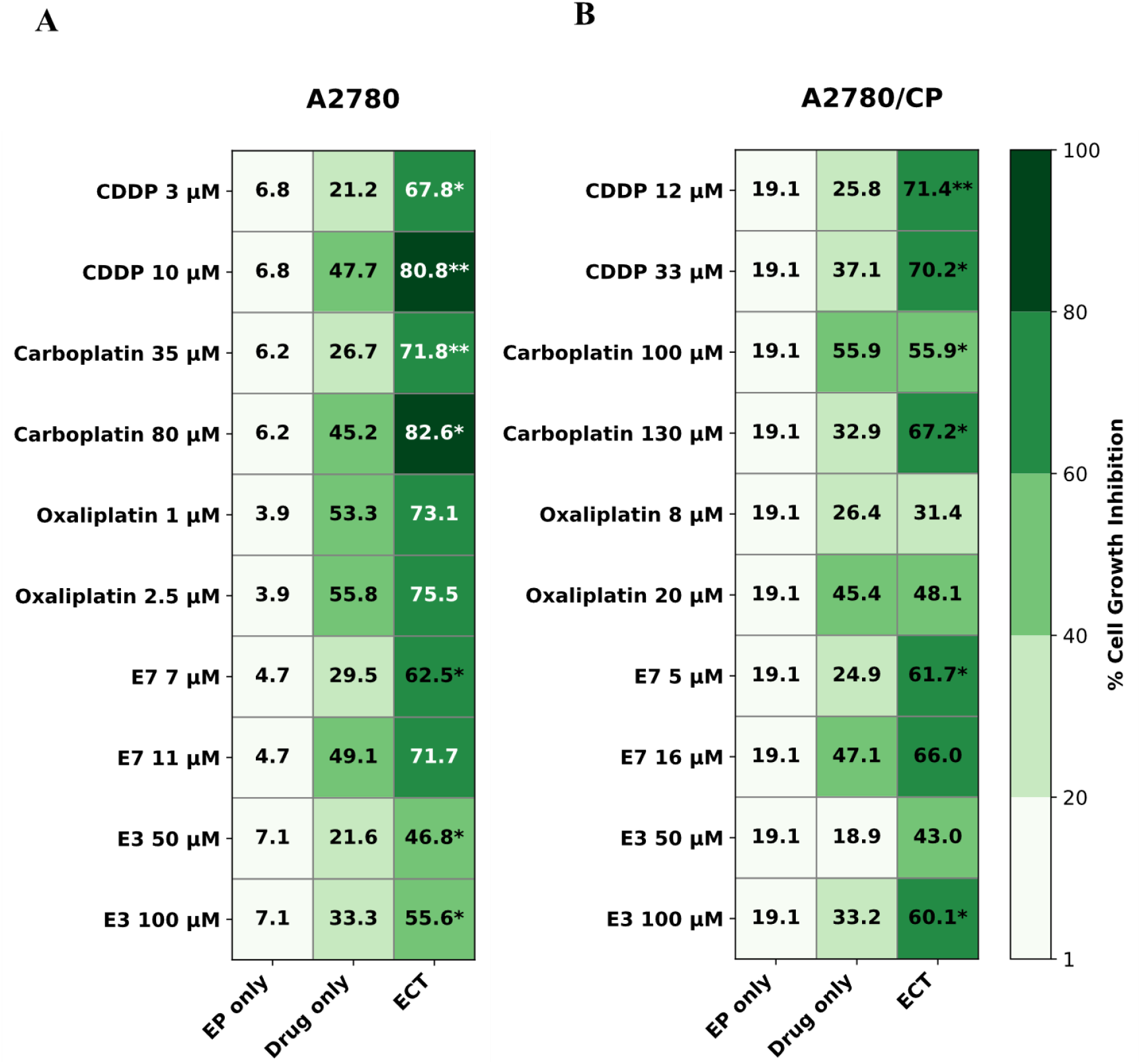
Heat-map representation of the cell growth inhibition by cDDP, CarboPt, OxaPt, **E7** and **E3** (IC20 and IC50) alone and in combination with 1100 V/cm electric pulses after 72h treatment in A2780 (A) and A2780/CP (B) cell lines. The numbers indicate the exact mean values of the inhibition data obtained for each treatment from at least three experiments. *P <0.05; **P <0.01, comparing the effect of the drug alone and in combination with EP. EP, electroporation. ECT, electrochemotherapy. As indicated by the colour scale on the right, the darker the colour, the greater the inhibition.

The cell growth inhibition effect of **E7** at the IC20 concentration increases from approximately 30% to 62% (p<0.05), while **E7** inhibition at the IC50 concentration increases from approximately 50% to ∼70%. Regarding **E3**, the cellular inhibition efficacy was almost doubled in EP conditions at both IC20 and IC50 concentrations. Both increases in cytotoxicity caused by the association with EP are significant at p<0.05 compared to the effect of the drug alone.

The same compounds tested on the A2780 cell line were also tested on the corresponding cisplatin-resistant line, A2780/CP.

**Figure 4B** depicts the increase of cDDP cytotoxicity by EP of about 45% and 30% at IC20 and IC50 concentrations, respectively. The IC20 concentration of CarboPt alone caused approximately 20% of cell growth inhibition and 55% in the presence of EP. Similar results in cell killing were also observed for the IC50 concentration. Both drugs at both concentrations were significantly potentiated by EP. Again, as observed in A2780 cells, even on the A2780/CP cell line, OxaPt appears to benefit less from the combination with EP, showing cellular inhibition following EP nearly the same as the drug alone. Regarding the two novel TS inhibitors, **E7** 5μM increased cell growth inhibition from approximately 25% to ~61% when combined with electric pulses and from approximately 47% to ~66% at the IC50 of 16 μM. EP also increases the cytotoxicity of **E3** by 20%, both at IC20 and IC50 concentrations (**Fig. 4B)**. In this cell line, EP significantly enhanced the efficacy of IC20 **E7** and IC50 **E3**, respectively.

Colorectal carcinoma cells, HCT116, were tested with OxaPt, 5-FU, **E7**, and **E3**. cDDP and CarboPt were not tested on these cells, since they are not primarily clinically used in the treatment of this type of tumor. OxaPt treatment at the IC50 3.5 µM in combination with EP provided significant increase of cell inhibition by about 35% (P<0.05), while only by 10% at 1.5 µM (**Fig. 5)**. As shown, ECT treatment doubles the inhibition of IC20 5-FU and increases cell growth inhibition of IC50 5-FU from ~ 50% to ~70% (P<0.05).

**Figure 5.**
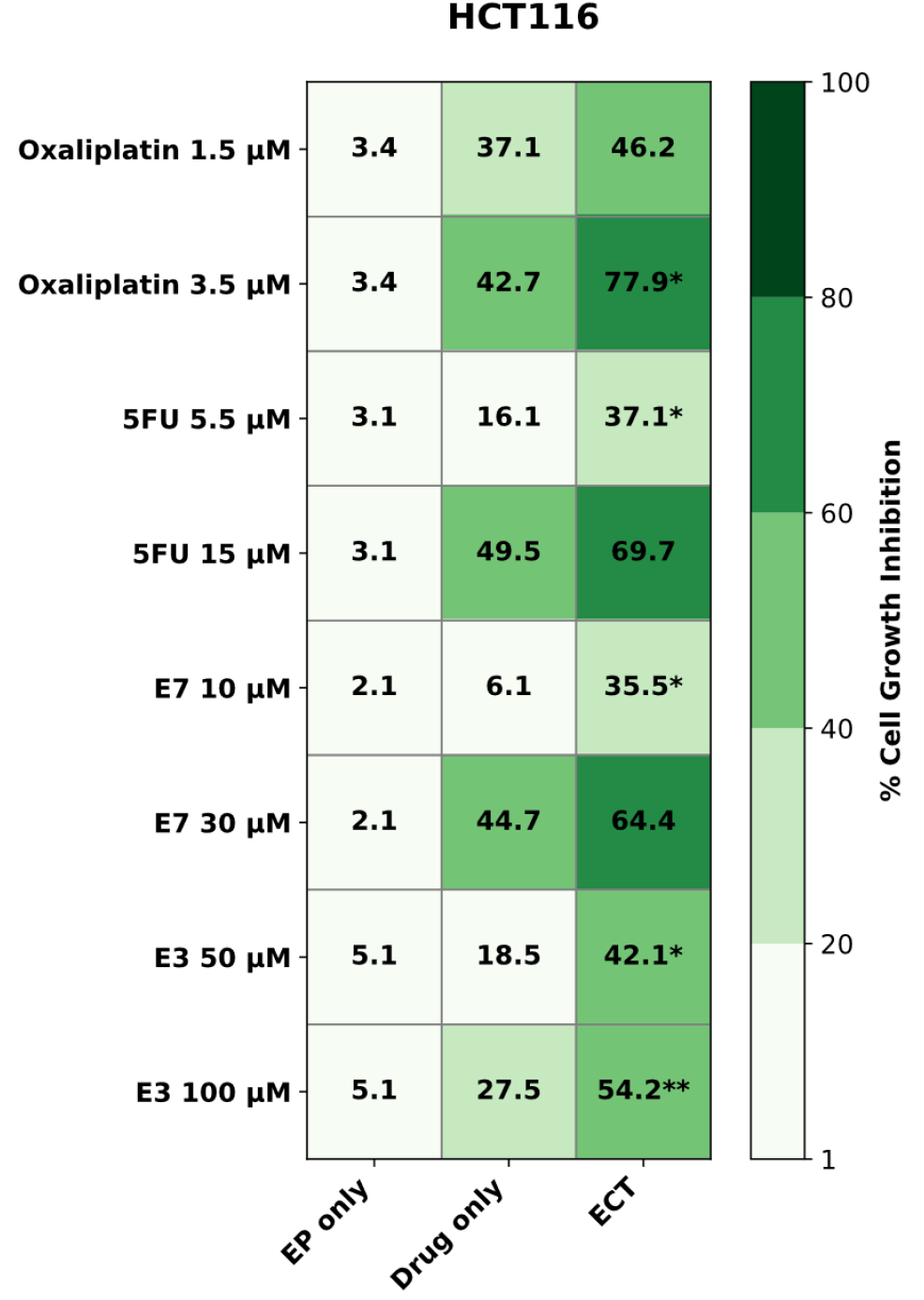
Heat-map representation of the cell growth inhibition by OxaPt, 5FU, **E7** and **E3** (IC20 and IC50) alone and in combination with 1100 V/cm electric pulses after 72h treatment in HCT116 cells. The numbers indicate the exact mean values of the inhibition data obtained for each treatment from at least three experiments. *P <0.05; **P <0.01, comparing the effect of the drug alone and in combination with EP. EP, electroporation. ECT, electrochemotherapy. As indicated by the colour scale on the right, the darker the colour, the greater the inhibition.

The cytotoxicity of 10 µM **E7** (IC20) increases from 6% alone to 35% in combination with EP (P<0.05), while 30 µM **E7** (IC50) enhanced cytotoxicity from 45% alone to 65% after EP.

**E3**, poorly lipophilic, also benefits from treatment with ECT, since its cell inhibition is doubled after treatment with ECT compared with treatment with the drug alone, both at IC20 and IC50 concentrations, P<0.05 and P<0.01, respectively.

### 3.4. Synergy analysis

To assess whether treatment with ECT really leads to an advantage over individual treatments with EP and drug alone, a synergy quotient (SQ) like analysis was performed. SQ is defined as the ratio between the inhibition obtained after treatment with ECT and the sum of the inhibitions of EP and the drug alone. Synergy occurs when the combined effect exceeds the effect of the individual factors, and the result is greater than 1. As shown in **Figure 6**, regarding treatments on the A2780 line, all drug-EP combinations have a synergistic effect (SQ>1.1) at both concentrations (IC20 and IC50).

**Figure 6.**
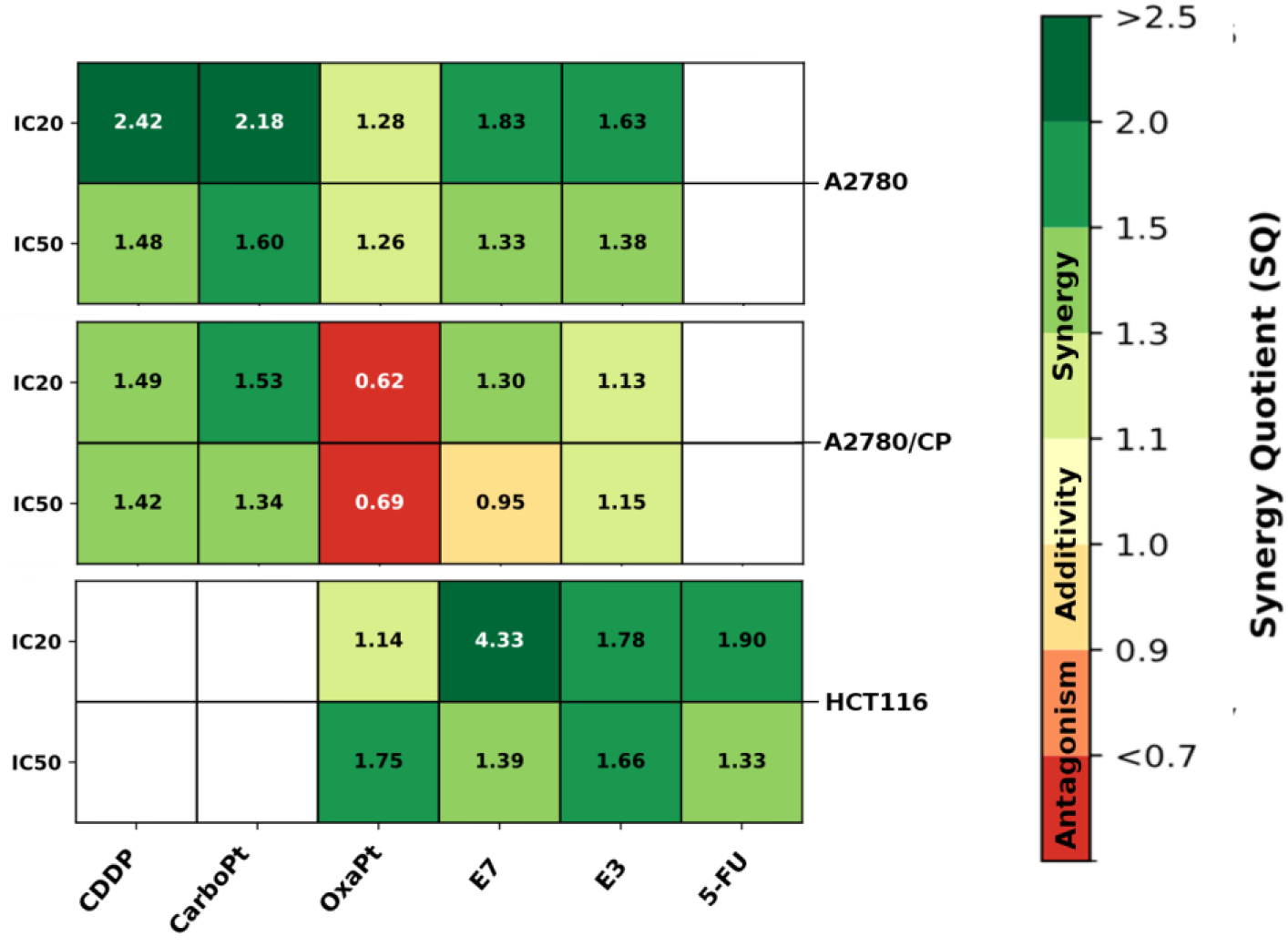
Heat-map representation of the Synergism Quotient (SQ) like analysis of the cytotoxicity of scheduled combinations of EP with the indicated tested compounds in A2780, A2780/CP and HCT116 cell lines. The numbers represent the mean of duplicate cell counts on two separate experiments and indicate the result of the net growth inhibitory effect of the combination of EP and drug divided by the sum of the net individual effects of drug and EP on growth inhibition. A quotient >1 indicates synergism (from light to dark green), while a quotient <1 or between 0.9 and 1.1 indicates an antagonistic (from orange to red) or additive effect (from white to yellow), respectively.

In A2780/CP cells, OxaPt did not take advantage of EP, showing antagonist values at both concentrations (SQ<0.9). IC50 **E7** showed an additive effect, while all the other drugs, including **E7** IC20, had a synergistic cell killing effect with EP.

The results on HCT116 cells show that all compounds tested in combination with EP have a synergistic effect, and therefore, there is a real advantage in treating these CRC cells with ECT. In particular, **E7** IC20 played with a particularly strong synergistic effect (SQ 4.3).

All the data of cell growth inhibition by single treatments and their combination with the resulting SQ values have been collected in **Table 1**.

**Table 1.** SQ values obtained from the various drug-EP combinations on the three cell lines. SQ>1.1 indicates a synergistic effect; 0.9<SQ<1.1 indicates an additive effect; SQ<0.9 indicates an antagonist effect.

| EP +<br>IC20<br>drugs | A2780 cells |  | A2780/CP cells |  | HCT116 cells |  |
| --- | --- | --- | --- | --- | --- | --- |
|  | % Growth inhibition | SQ | % Growth inhibition | SQ | % Growth inhibition | SQ |
| cDDP | 67.8/6.8+21.2 | 2.42 | 71.4 / (22.1+25.8) | 1.49 | - | - |
| CarboPt | 71.8/6.2+26.7 | 2.18 | 55.9 / (17.9+18.5) | 1.53 | - | - |
| OxaPt | 73.1/3.9+53.3 | **1.28** | 31.4 / (24.1+26.4) | **0.62** | 46.2/3.4+37.1 | **1.14** |
| E7 | 62.5/4.7+29.5 | **1.83** | 61.7 / (22.5+24.9) | **1.3** | 35.5/2.1+6.1 | **4.33** |
| E3 | 46.8/7.1+21.6 | **1.63** | 43.0 / (19.2+18.9) | **1.13** | 42.1/5.1+18.5 | **1.78** |
| 5-FU | - | - | - | - | 37.1/3.1+16.4 | **1.90** |
| **EP +**<br>**IC50**<br>**drugs** |  |  |  |  |  |  |
| cDDP | 80.1/6.8+47.7 | **1.48** | 70.2 / (22.1+37.1) | **1.42** | - | - |
| CarboPt | 82.6/6.2+45.2 | **1.60** | 67.2 / (17.9+32.9) | **1.34** | - | - |
| OxaPt | 75.5/3.9+55.8 | **1.26** | 48.1 / (24.1+45.4) | **0.69** | 77.9/3.4+41.1 | **1.75** |
| E7 | 71.7/4.7+49.1 | **1.33** | 66.0 / (22.5+47.1) | **0.95** | 64.9/2.1+44.7 | **1.39** |
| E3 | 55.6/7.1+33.3 | **1.38** | 60.1 / (19.2+33.2) | **1.15** | 54.2/5.1+27.6 | **1.66** |
| 5-FU | - | - | - | - | 69.7/3.1+49.5 | **1.33** |

### 3.5. ECT with drug combinations

Subsequently, it was decided to test combinations of two drugs with EP, on A2780 and HCT116 cells. All compounds were tested at their IC20 concentration. **E7**, the most active Ddis compound, was tested with platinum derivatives (cDDP, CarboPt, OxaPt) on A2780 cells, before and after treatment with EP. On HCT116 cells, **E7** was tested with CarboPt, OxaPt and 5-FU.

Figure 7. shows the cellular inhibition by treatments performed on the two different tumor cell lines. EP increases the cytotoxicity of all combinations tested. The cytotoxicity of **E7**+CarboPt is significantly enhanced by 35% compared to the counterpart without EP. For the other combinations, **E7**+cDDP and **E7**+OxaPt, the addition of EP increases cellular inhibition by 15%. In HCT116 cells, all two drug treatments gained benefit from EP, and more significantly in **E7**+cDDP, increasing cellular inhibition.

## 4. Discussion

Many human cancers, including ovarian and colorectal cancers, have been found to develop resistance to multiple chemotherapeutic drugs, also referred to as multi-drug resistance (MDR), which limits chemotherapy efficacy for cancer treatment. The MDR effect is an insensitivity of cancer cells not only to previously administered drugs but also to many other drugs with different chemical structures and mechanisms of action (Ullah, 2008). MDR is usually associated with reduced tumoral drug accumulation and increased removal of the drug from the tumor. The latter mechanism is due to the overexpression of ATP-binding cassette (ABC) transporters such as P-glycoproteins (P-gp), multidrug-resistance protein (MRP), and breast cancer resistance protein (BCRP) on the surface of cancer cells (Teodori et al., 2006; Lee, 2004). ABC transporters contribute to both intrinsic and acquired drug resistance in colon and ovarian cancer (Baekelandt et al., 2000), where this transmembrane protein mediates cellular efflux of many chemotherapeutic agents clinically used against this cancer, including cisplatin (Ortiz et al., 2022). P-gp expression is also inducible by chemotherapeutic agents in colon cell lines, including HCT116 and HT29 (Ding et al., 2010). Besides, P-gp, MRP1, BCRP and several other ABC transporters could also be induced by anticancer drugs targeting the folate cycle enzymes in colon cancer cells and play roles in drug resistance. For instance, ABCB5 expression was substantially increased in clinical colorectal cancers after 5-FU-based chemotherapy and contributed to the development of resistance to 5-FU (Wilson et al., 2011).

**Figure 7.**
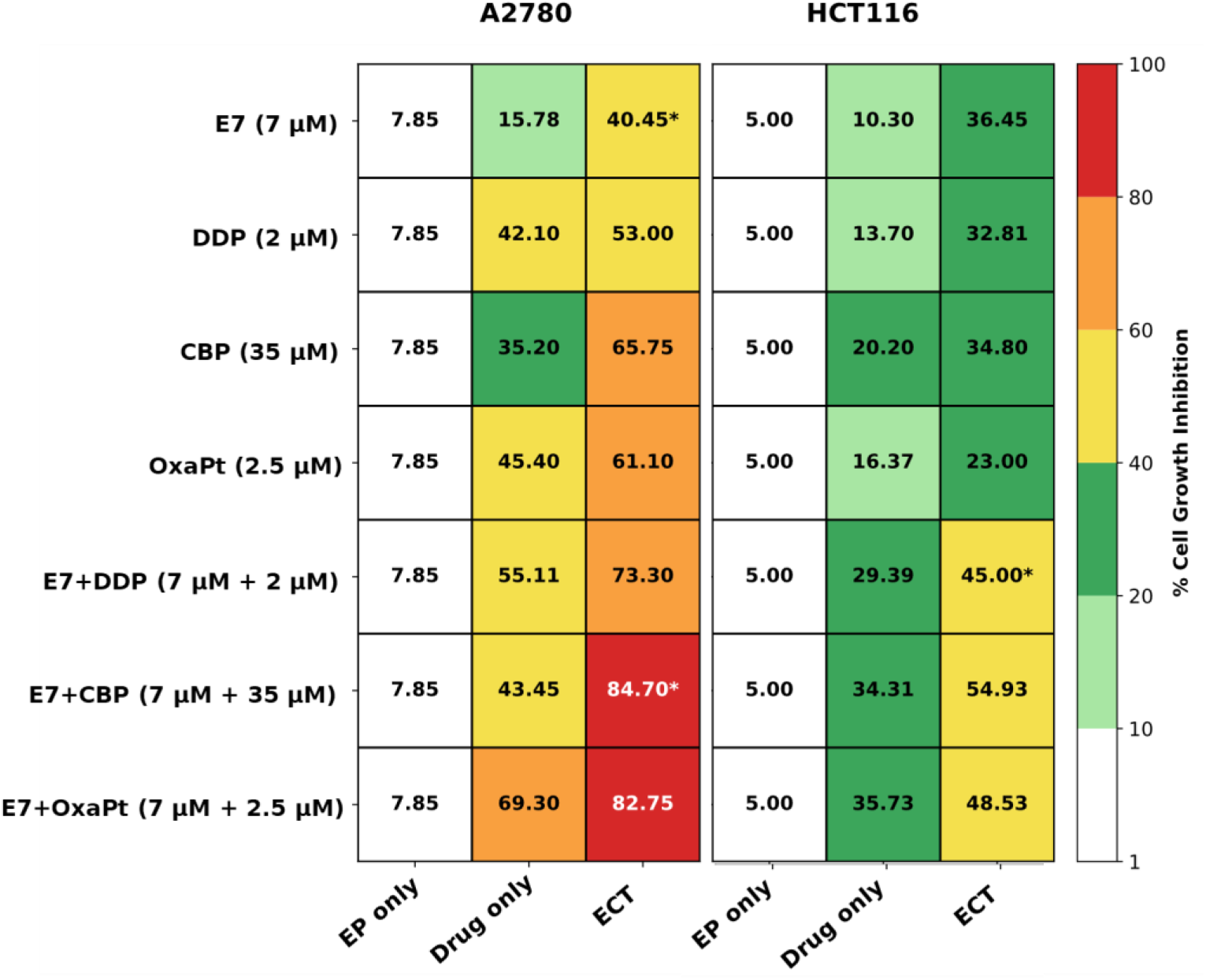
Heat-map representation of A2780 and HCT116 cell growth inhibition by Pt-drugs and **E7** alone or in combination at IC20 ± 1100 V/cm electric pulses after 72h treatment. The numbers indicate the exact mean values of the inhibition data obtained for each treatment from at least two experiments. *P <0.05, comparing the effect of the drug alone and in combination with EP. EP, electroporation. ECT, electrochemotherapy. As shown by the colour scale on the right, the increased growth inhibition is indicated by the intensity of the different colours.

Among the novel strategies to overcome resistance mechanisms for TS and TS-pathway-targeted drugs, we have recently developed the TS-dimer disrupters (Ddis), compounds that bind at the monomer-monomer interface and shift the dimerization equilibrium of both the recombinant and the intracellular protein toward the inactive monomers (Costantino et al., 2022). Among them, one molecule in particular, **E7**, shows a superior anticancer profile to 5-FU in a mouse model of human pancreatic cancer, and it inhibits colorectal and ovarian cancer cell growth. Other Ddis TS inhibitors, such as **E3**, have shown interesting inhibitory activity at the enzyme level (Kd=7 μM), but limited cellular growth inhibition activity, thus suggesting that the moderate effect at the cellular level may be due to the low trans-membrane internalization process. This event is competitive with the interaction of the compounds with the intracellular target and may reduce the Ddis-TS availability and the effective amount administered to develop their inhibitory activity.

Recently, by using specific MDR efflux pump inhibitors to prevent the Ddis efflux out of the cells, we have favoured their internalization, enhanced their accumulation and cytotoxicity (Marverti et al., 2024). The results demonstrated that these novel TS inhibitors could be substrates for the extrusion protein pumps whose inhibition makes the Ddis more effective, including both the most active **E7** and the least active **E3**.

As a second strategy to increase Ddis cellular accumulation, we enhanced drug uptake by transfection of the Ddis in an alternative exploitation of a specific peptide delivery system, the SAINT-Protein (Synvolux Products & Therapeutics, Leiden, The Netherlands), a non-liposomal lipid reagent.

The Ddis compounds, and also 5FU, benefited from combination with SAINT-Protein, especially compounds like **E3**, that inhibit the recombinant TS activity *in vitro*, but show their cytotoxic efficacy limited by the obstacle of biological membranes (Marverti et al., 2024).

In this study, we focused on overcoming drug uptake limitations by forcing drug internalization by means of electroporation (EP), a physical phenomenon that occurs when a sufficiently large electric field is applied to a tissue for an adequate duration. In this case, a transmembrane voltage is induced across the cell plasma membrane, causing local deficiencies and establishing pores that make it permeable to molecules that otherwise could not transfer into it, such as water, charged and larger molecules. The pore is formed rapidly and disappears within a few seconds to several minutes after exposure to the electric field. The duration of the pulses and the electric field intensity determine whether the structural changes in the cell membrane are reversible, allowing cells to survive, or irreversible, leading to cell death because of the loss of homeostasis.

EP permeates mammalian cell membranes carrying out high-frequency electrical pulses in saline buffer solution or culture medium with pulses of 400-1500 V/cm lasting 10-40 milliseconds that cause the transmembrane voltage (ΔU_(t)_) to increase from the physiological value of 0.1 up to 0.5-1 V (Kumar et al., 2019).

Three main steps in the electroporation mechanism can be identified: induction (application of the electric field), expansion (the electric impulse causes pores to form in the membrane, which expand until the impulse ceases), and resealing (repairing the membrane). If the cell is subjected only temporarily, for a short period of time, to the electric field, it will be able to repair the membrane properly: this process is called reversible EP. On the other hand, in the irreversible EP, the cell membrane loses its integrity, and the cell goes into apoptosis (Tasu et al., 2022).

To date, there is a therapeutic approach for liver, pancreatic, kidney and prostate tumors based on irreversible electroporation (IRE): administration occurs percutaneously via imaging or during surgery. IRE has emerged as an emerging technique for the treatment of early-stage tumors. Reversible electroporation finds application as a method for transferring material (such as DNA, plasmids, drugs) inside cells. EP facilitates the passage of therapeutic agents across the plasma membrane, increasing their efficacy and has been much studied in oncology.

Electrochemotherapy (ECT) combines reversible EP with the administration of chemotherapeutic agents with low cytotoxicity, also allowing for a more localized distribution of the drug, reducing any side effects (Cucu et al., 2021; Cemazar and Sersa, 2019; Bieżuńska-Kusiak et al., 2023).

Different cells respond differently to electrical pulses: variations in electroporation efficiency are due to differences in membrane recovery at the end of the pulse. In addition, cell diameter, pulse length, electric field strength and direction are also parameters that affect electroporation efficiency.

Therefore, we have applied ECT to two ovarian carcinoma cell lines, one cDDP-sensitive and the other cDDP–resistant, and one colorectal cell line innately poorly sensitive to 5FU, to verify whether the positive effect of the electric field on the action of the drugs chosen could have a valid application to two different and aggressive tumor types.

Several studies have been conducted to evaluate the effect of ECT with cDDP on ovarian cancer cell lines, showing that EP promotes the distribution and cytotoxicity of cisplatin both in human ovarian cancer cells (IGROV1) and in its cisplatin-resistant subclone (IGROV/DDP) (Perrone et al., 2021; Łapińska et al., 2022; Saczko et al., 2016).

Studies have also been conducted on cisplatin-resistant ovarian cancer cell lines (OvBH-1 and SKOV-3); to find a solution to the drug resistance that these tumors show after treatment with platinum derivatives, some researchers have treated these cells with a combination of EP and 5-FU. They have observed that treatment with ECT has greater efficacy than chemotherapy treatment alone, and EP also increases the efficacy of cDDP, despite these cells being resistant to it, because cellular uptake is increased (Saczko et al 2014).

According to these studies, all treatments we tested with the ECT protocol were shown to cause enhanced cellular growth inhibition compared to their counterparts without EP, except OxaPt. In particular, cDDP and CarboPt show a synergistic effect in combination with EP at both concentrations on ovarian cells. The greatest results were obtained on the A2780 cell line, in which treatments with cDDP, CarboPt and also **E3** at the IC20 concentration demonstrated a significant increase in cytotoxicity. cDDP was also found to be highly active in the A2780/CP cell line, which is resistant to this drug. Thus, according to other reports (Perrone et al., 2021; Łapińska et al., 2022) this result reinforces the concept that EP, facilitating the uptake of cDDP, may overcome resistance mechanisms mainly based on accumulation limits and allow for greater drug effectiveness.

The more rapid and higher intracellular concentration of these drugs reached with EP allows them to bypass the limitations of the uptake mechanism as well as of extrusion pumps characteristics of resistant cells, including the binding with inactivating intracellular substances such as GSH and metallothioneins, often overexpressed in resistant cell lines. In this way, a higher number of drug molecules can reach the main target, the DNA, and inhibit its metabolism by forming more DNA adducts and causing DNA damage, counteracting resistance. OxaPt is the drug that benefited less from electroporation, probably because its lipophilic structure already guarantees an adequate cellular uptake, and treatment with EP does not add a significant pharmacokinetic advantage.

On the other hand, according to Saczko *et al*. (Saczko et al. 2014), our experiments also show that EP increases the cytotoxicity of 5-FU, a traditional TS inhibitor, even on HCT116 cells, innately 5FU-resistant. 5FU and its derivatives, including the more active FdUMP, can be substrates of the ATP binding cassettes, in particular, ABCC5, ABCC11 and ABCG2 (Sethy and Kundu, 2011), thus, it is likely that EP allows more 5FU to enter cells and to form more FdUMP than that extruded by ABC pumps.

The two new TS-dissociative inhibitor compounds also showed increased cytotoxic activity after treatment with EP. The action of **E3** is limited by reduced uptake across the plasma membrane, but under EP conditions, it increased its cytotoxicity due to increased intracellular accumulation, producing a synergistic effect on two lines and an additive on the third cell line. **E7** is the one shown to have greater inhibitory activity alone, but it increased further more by EP; in fact, the IC20 and IC50 values of **E7** are lower than those of **E3**, because it has better cellular uptake that allows the use of lower concentrations of drug to achieve the cytotoxic effect.

Like transfection with SAINT-Protein (Marverti et al., 2024), the Ddis compounds benefited from combination with EP, whose role as a facilitator in overcoming biological barriers was especially useful for those compounds that, like **E3**, despite inhibiting the activity of the recombinant *h*TS enzyme *in vitro*, show limited cytotoxicity since they alone cannot reach an effective intracellular concentration. Thanks to EP, they can effectively kill even resistant tumor cells at the IC20 concentration; and even in this case, the increased cytotoxicity may likely correlate with enhanced cell cycle perturbation and apoptosis, and the reduction of *h*TS protein levels (Marverti et al., 2024). Notably, we demonstrate that EP significantly increases the efficacy of drug combinations in both OC and CRC cells. EP significantly (P<0.05) enhanced cell growth inhibition when **E7** was combined with platinum-based drugs, specifically CarboPt in OC and OxaPt in CRC. Interestingly, this combination treatment potentiated the effects of OxaPt, even though its action as a single agent was not enhanced by EP. Under these conditions, the intracellular concentration of **E7** achieved via EP likely inhibits TS to such an extent that it impairs the cell’s ability to repair DNA damage induced by platinum drugs.

Consequently, EP proved to be an effective method for permeabilizing the cell membrane, allowing drugs, particularly those with lower lipophilicity to reach intracellular concentrations necessary to engage their targets, whether administered alone or in combination. This study indicates that EP can effectively enhance the potency of both traditional chemotherapeutics and novel allosteric TS inhibitors, offering a potential strategy for treating OC and CRC cells resistant to standard therapies.

## Author Contributions

Conceptualization, G.M., M.P.C. and D.D.; methodology, G.M., A.B., G.Me., D.A., software, G.M., D.A.; validation M.G., M.P.C. and D.D.; formal analysis, M.G. and D.D.; investigation, G.M., M.P.C. and D.D.; resources; data curation, G.M., M.P.C. and D.D.; writing—original draft preparation, G.M., M.P.C. and D.D.; writing—review and editing, G.M., D.D. and M.P.C.; supervision, G.M., D.D., A.V., and M.P.C.; project administration, M.P.C.; funding acquisition, M.P.C. All authors have read and agreed to the published version of the manuscript.

## Acknowledgments

We thank the IGEA Electroporation (Carpi, Italy) for making the instrument Genedrive electroporator available for the experiments reported in the manuscript.

